# Screening a library of FDA-approved and bioactive compounds for antiviral activity against SARS-CoV-2

**DOI:** 10.1101/2020.12.30.424862

**Authors:** Scott B. Biering, Erik Van Dis, Eddie Wehri, Livia H. Yamashiro, Xammy Nguyenla, Claire Dugast-Darzacq, Thomas G.W. Graham, Julien R. Stroumza, Guillaume R. Golovkine, Allison W. Roberts, Daniel M. Fines, Jessica N. Spradlin, Carl C. Ward, Teena Bajaj, Dustin Dovala, Ursula Schulze Gahmen, Ruchika Bajaj, Douglas M. Fox, Melanie Ott, Niren Murthy, Daniel K. Nomura, Julia Schaletzky, Sarah A. Stanley

**Affiliations:** School of Public Health, Division of Infectious Diseases and Vaccinology, University of California, Berkeley, Berkeley, California, 94720, USA; Department of Molecular and Cell Biology, Division of Immunology and Pathogenesis, University of California, Berkeley, Berkeley, California, 94720, USA; The Henry Wheeler Center for Emerging and Neglected Diseases, 344 Li Ka Shing, Berkeley, California, 94720, USA; Department of Molecular and Cell Biology, Division of Biochemistry, Biophysics and Structural Biology, University of California, Berkeley, Berkeley, California 94720, USA; Departments of Chemistry, Molecular and Cell Biology, and Nutritional Sciences and Toxicology, University of California, Berkeley; Department of Bioengineering, University of California, Berkeley, Berkeley, CA 94720, USA; Novartis Institutes for BioMedical Research, Emeryville, CA, 94608, USA; QBI Coronavirus Research Group Structural Biology Consortium, University of California, San Francisco, CA 94158, USA; Department of Bioengineering and Therapeutic Sciences, University of California, San Francisco, San Francisco, CA 94158, USA; Department of Medicine; Medical Scientist Training Program; Biomedical Sciences Graduate Program, University of California, San Francisco, San Francisco, CA 94143, USA; J. David Gladstone Institutes, San Francisco, CA 94158, USA; Innovative Genomics Institute (IGI), 2151 Berkeley Way, CA 94704, USA

## Abstract

Severe acute respiratory syndrome coronavirus 2 (SARS-CoV-2), the causative agent of coronavirus disease 2019 (COVID-19), has emerged as a major global health threat. The COVID-19 pandemic has resulted in over 80 million cases and 1.7 million deaths to date while the number of cases continues to rise. With limited therapeutic options, the identification of safe and effective therapeutics is urgently needed. The repurposing of known clinical compounds holds the potential for rapid identification of drugs effective against SARS-CoV-2. Here we utilized a library of FDA-approved and well-studied preclinical and clinical compounds to screen for antivirals against SARS-CoV-2 in human pulmonary epithelial cells. We identified 13 compounds that exhibit potent antiviral activity across multiple orthogonal assays. Hits include known antivirals, compounds with anti-inflammatory activity, and compounds targeting host pathways such as kinases and proteases critical for SARS-CoV-2 replication. We identified seven compounds not previously reported to have activity against SARS-CoV-2, including B02, a human RAD51 inhibitor. We further demonstrated that B02 exhibits synergy with remdesivir, the only antiviral approved by the FDA to treat COVID-19, highlighting the potential for combination therapy. Taken together, our comparative compound screening strategy highlights the potential of drug repurposing screens to identify novel starting points for development of effective antiviral mono- or combination therapies to treat COVID-19.

## Introduction

Zoonotic viruses pose a great public health challenge due to the unpredictable nature of an outbreak, the potential to impact an immune-naïve population, and a lack of therapeutic options (*1, 2*). Coronavirus disease 2019 (COVID-19) is caused by the emergence of the severe acute respiratory syndrome coronavirus 2 (SARS-CoV-2), a member of the *Coronaviridae* family within the *Betacoronavirus* genus (*3–5*). The *Betacoronavirus* genus contains several seasonal human pathogens that cause mild disease, as well as the recently emerged severe acute respiratory syndrome coronavirus (SARS-CoV) and Middle East respiratory syndrome coronavirus (MERS-CoV) (*4*). SARS-CoV-2, SARS-CoV, and MERS-CoV cause severe disease in humans. The increased magnitude of the current pandemic relative to SARS-CoV and MERS-CoV may be explained by increased human-human transmissibility from frequent asymptomatic shedding of this virus (*6–9*). Severe cases of COVID-19 are associated with acute respiratory distress syndrome (ARDS) triggered by an inflammatory response resulting in tissue damage and fluid accumulation in the lungs (*10–12*). Currently there are limited options to treat patients suffering from severe COVID-19. The sole FDA-approved antiviral compound for COVID-19 treatment is remdesivir, but clinical efficacy is modest and no conclusive effect on patient mortality has been found (*13–16*). Although remdesivir exhibits strong *in vitro* efficacy against SARS-CoV-2 infection, its low exposure in target lung tissue, dose-limiting kidney and liver toxicity, and the need for intravenous administration make early and effective clinical treatment difficult (*17–20*). Consequently, the rapid discovery and development of additional therapeutics is vital.

Repurposing well-studied preclinical, clinical, and approved compounds holds the greatest potential to swiftly move a drug candidate from the bench to the clinic. An FDA-approved compound library (TargetMol L4200) and a bioactive compound library (TargetMol L4000) are two collections of well-studied compounds, many of which already possess extensive human safety data. Here, we screened both libraries to identify compounds that inhibit SARS-CoV-2 replication in host cells. We report the identification of 13 compounds effective against SARS-CoV-2, 7 of which are previously unreported. Of note, a host-directed compound, the cyclin-dependent kinase (CDK) inhibitor dinaciclib, was determined to be more potent than remdesivir in limiting viral replication in human lung epithelial cells. Additional hit compounds include: i) known antivirals predicted to target SARS-CoV-2 viral factors directly; ii) host protein kinase and protease inhibitors; and iii) and anti-inflammatory agents. Further, we identify antiviral synergy between remdesivir and the RAD51 inhibitor B02, opening the possibility of clinical combination therapy. Taken together, our results identify several novel starting points for COVID-19 drug development which hold the potential to alleviate the current global pandemic.

## Results

### Identification of SARS-CoV-2 permissive cell lines for drug screening

To design a screen for identification of SARS-CoV-2 antiviral compounds, we first tested a panel of 16 cell lines to identify cells capable of sustaining robust SARS-CoV-2 replication. These cell lines included human pulmonary epithelial and endothelial cells, keratinocytes, hepatocytes, and primate cells, among others (**Figure 1 and Figure S1**). Multi-step growth curves of SARS-CoV-2 in each of these cell lines revealed a range of permissiveness to viral infection (summarized in **Table 1)**. We found distinct cell infection patterns including highly permissive for viral replication **(Figure 1A-D)** with and without significant cytopathic effect (CPE) **(Table 1)**, mildly permissive with no CPE **(Figure S1, Table 1)**, as well as non-permissive defined by lack of detectable infectious virus measured by a median tissue culture infectious dose assay (TCID50) **(Figure S1, Table 1)**. We selected the human pulmonary epithelial cell line Calu-3 and the African Green Monkey kidney cell line Vero-E6 for conducting the primary screens, as these cell types supported high levels of infection with dramatic CPE, ideal features for drug screening **(Figure 1A and 1B)**. We further reasoned that selection of two distinct cell lines would control for cellspecific effects of a given compound.

**Figure 1.**
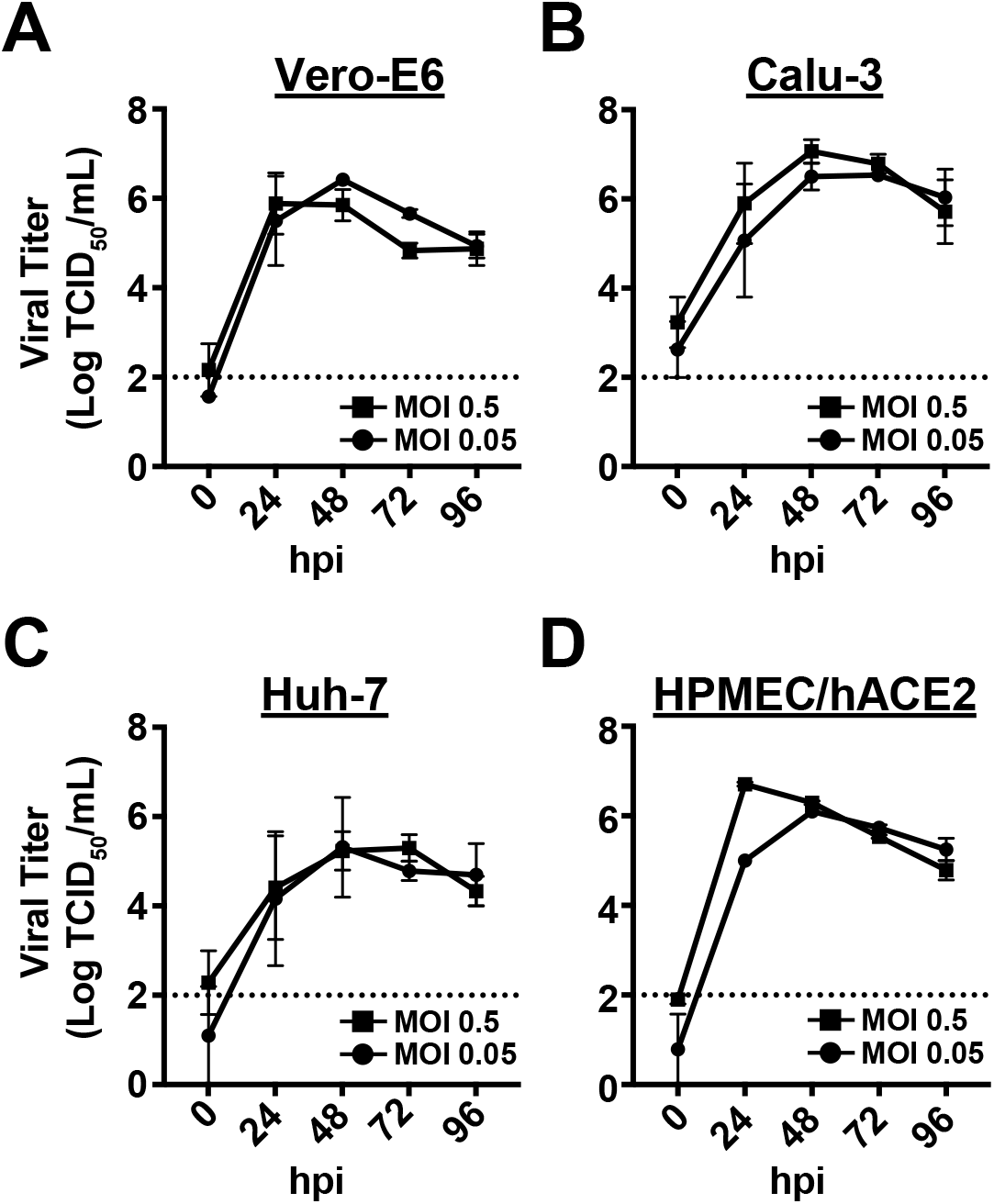
Permissive cell lines to SARS-CoV2 infection. **(A)** Vero-E6, **(B)** Calu-3, **(C)** Huh-7 and **(D)** HPMEC/hACE2 cells were seeded in 24-well plates and infected with SARS-CoV-2 at MOI 0.5 or 0.05 at 37°C and 5% C0_2_ for 30 minutes. Viral inoculum was then removed, cells were washed once in 1x PBS, and 1 ml of regular media was replaced. At the indicated time points (hours post-infection, hpi), plates were freeze/thawed and viral titers from whole cell lysates were analyzed by TCID50 assay. Dashed line represents limit of detection of the assay. Data represent mean ± SEM for n = 2 independent experiments.

**Table 1.**

| Cell Line | Species | Origin | Permissive | Cytopathic Effect (CPE) |
| --- | --- | --- | --- | --- |
| Vero-E6 | African Green Monkey | Kidney Epithelial Cell | yes | +++ |
| Calu-3 | Human | Lung Epithelial Cell, adenocarcinoma | yes | +++ |
| HPMEC/hACE2 | Human | Lung Microvascular Endothelial Cell, primary | yes | +++ |
| A549/hACE2 | Human | Lung Epithelial Cell, carcinoma | yes | ++ |
| Huh-7.5.1 | Human | Hepatocytes, carcinoma | yes | ++ |
| Huh-7 | Human | Hepatocytes, carcinoma | yes | + |
| Caco-2 | Human | Colon Epithelial Cell, adenocarcinoma | yes | - |
| NCI-H1437 | Human | Patient Lung Cell derived from Metastatic Site | yes | - |
| LNCaP | Human | Prostate Epithelial Cell, carcinoma | yes | - |
| HaCaT | Human | Keratinocyte, transformed | yes | - |
| HCT-116 | Human | Colon Epithelial Cell, colorectal carcinoma | no | - |
| A549 | Human | Lung Epithelial Cell, carcinoma | no | - |
| HBEC-30KT | Human | Bronchial Epithelial Cell, immortalized | no | - |
| BEAS-2B | Human | Bronchial Epithelial Cell, transformed | no | - |
| RD | Human | Derived from the muscle, rhabdomyosarcoma | no | - |
| HPMEC | Human | Lung Microvascular Endothelial Cell, primary | no | - |

### A cytopathic effect-based screening platform to identify SARS-CoV-2 antiviral compounds from the FDA-approved and bioactive compound libraries

We screened a library of 1,200 FDA approved compounds (FDA-approved library) and a library of 2,834 pre-clinical compounds and 1,336 compounds with human clinical data (bioactive library, **Figure S2A)**. For the primary screen, we developed a CPE-based screening assay (CellTiter-Glo, CTG) which uses ATP released from viable cells to drive a luciferase reporter (RLU). This assay identifies drugs that are potent enough to inhibit SARS-CoV-2-mediated cell death, thus selecting for compounds whose antiviral effects are present over several viral lifecycles. Cells are treated with compound immediately prior to infection with SARS-CoV-2 and incubated until complete CPE is observed in infected vehicle treated wells, at which time cell viability is determined using the CTG assay **(Figure 2A)**. We used both Calu-3 and Vero-E6 cells to identify possible cell type dependent effects of each compound. We observed significant separation in signal between infected and uninfected wells in Vero-E6 cells at day 3 postinfection **(Figure 2B,** Z’=0.47) and in Calu-3 cells at day 4 post-infection **(Figure 2C**, Z’=0.43), demonstrating that the assay was suitable for high-throughput screening. Remdesivir served as a positive control in this assay, with EC50 values of 0.7-3μM **(Figure S2B, S2C)**. For primary screens, three identical 384-well daughter plates were generated at a final compound concentration of 40 μM. Two plates served as technical replicates that were infected with SARS-CoV-2, while the third functioned as an uninfected cell cytotoxicity control (**Figure 2A**).

**Figure 2.**
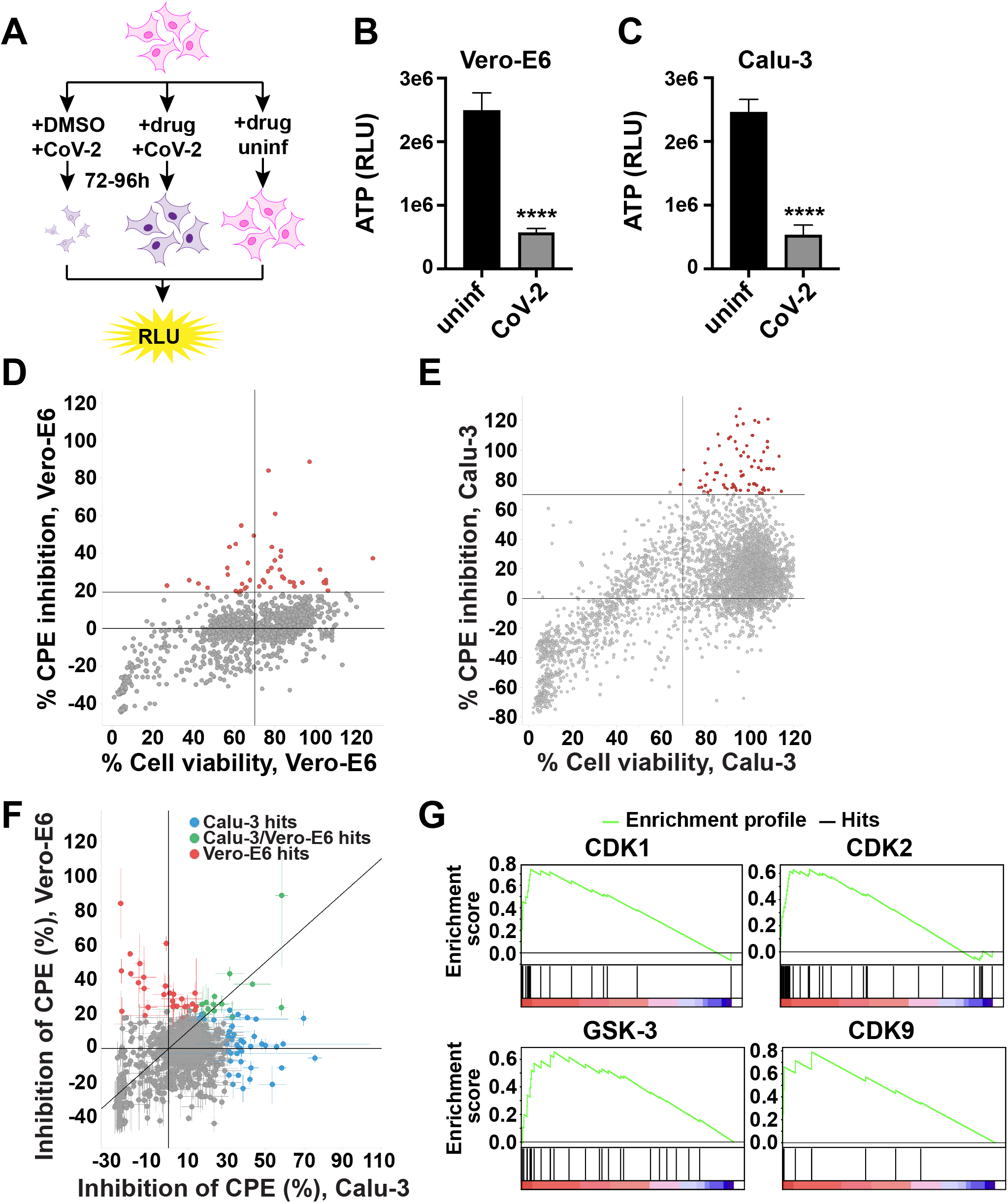
Screening SARS-CoV-2 antiviral activity using the FDA-approved and bioactive compound libraries. **(A)** Assay scheme: Cells are treated with DMSO (left panel) or drug (middle and right panels), infected with SARS-CoV-2 or left uninfected (right panel) and incubated for 72-96h to observe cytopathic effect (CPE). CPE is measured by CTG assay, quantifying ATP content in viable cells using luminescence (RLU). The right panel shows the cytotoxicity control, treating cells with drugs but without virus. **(B-C)** Average luminescence is shown for **(B)** Vero-E6 at 72h or **(C)** Calu-3 cells at 96h post-infection. **(D)** Screen of FDA-approved and bioactive compound libraries on Vero-E6 cells with inhibition of CPE (%) on the y-axis and cell viability (%) on the x-axis normalized to DMSO-treated wells. Red: high priority hits with a cutoff of >20% inhibition of CPE and >70% cell viability. **(E)** As in (D), but on Calu-3 cells, with a cutoff of >70% inhibition of CPE and >70% cell viability. **(F)** Combination of inhibition of CPE (%) on Vero-E6 (y-axis) from (D) and Calu-3 (x-axis) from (E). **(G)** Gene set enrichment analysis. Distribution of the enrichment score (green line) across compounds annotated to molecular targets (vertical black lines). CDK1, CDK2, GSK-3 p < 0.001, CDK9 p = 0.0015. False discovery rate (FDR) q < 0.05. Data represent mean ± SEM for n = 24 technical replicates (B, C) or n = 2 technical replicates (F). ****: p < 0.0001.

To enable comparative analysis, we ran two parallel screens using the FDA-approved compound library on both Vero-E6 and Calu-3 cells. These screens revealed numerous compounds that significantly inhibited CPE in both cell types without causing significant cytotoxicity in uninfected cells (**Figure 2D-F**). Intriguingly, although some compounds displayed overlapping antiviral activity in both cell lines, the majority displayed cell specific antiviral activity (**Figure 2F**). These observations could be explained by species-specific or cell type-specific mechanisms of action of a given compound, or by differences in viral replication between the two cell types. Because we observed cell type-specific effects of compounds, we proceeded to screen the remaining compounds from the bioactive library using Calu-3 human pulmonary epithelial cells only, which is more physiologically relevant to human infection. Selecting a hit signal cutoff of >1.5 and >2.1 standard deviations from the library mean for the FDA approved and bioactive library, respectively, our screens identified a total of 140 unconfirmed compounds that significantly inhibit SARS-CoV-2-mediated CPE **(Figure 2E and 2F)**. Best hits were selected for dose response assays based both on these primary screen data and an assessment of their suitability as potential COVID-19 therapeutics and prioritized for follow-up validation experiments using an experimental pipeline designed to narrow down our list of candidates to only the most promising compounds.

Leveraging the fact that compounds in the FDA-approved and bioactive libraries are well characterized with many drug targets previously defined, we conducted a gene set enrichment analysis (GSEA) of our candidate compounds to shed light on host-pathways important for SARS-CoV-2 replication. We found significant enrichment of compounds targeting host cyclin-dependent kinases (CDK1, CDK2, and CDK9), GSK-3β, C-RAF, and JNK1-3, suggesting that ubiquitous host pathways such as cell-cycle progression, MAPK-signaling, as well as GSK-3-signaling are critical for SARS-CoV-2 infection. This agrees with a previously published analysis based on phosphoproteomics of cells infected with SARS-CoV-2 (*21*) **(Figure 2G)**. Thus, our screen suggests these cellular pathways may contain druggable targets for inhibition SARS-CoV-2 infection of human lung cells.

To validate the *in vitro* potency of drug candidates revealed in our screen, we determined the half-maximal effective concentrations (EC50) of each compound by conducting dose response curves in the cell line in which compounds were identified using our optimized CTG system as a readout **(Figure S2C)**. Confirmation rates in the secondary dose response screen were 72% for the Calu-3 and 67% for the Vero-E6 screen, highlighting assay reproducibility and suitability for hit identification **(Figure 3, Figure S3)**. Our results identified 17 compounds with EC50 values below 10 μM, including 6 below 5 μM, without significant cytotoxicity. Data for the most potent compounds in Calu-3 cells are shown in **Figure 3** and **Figure S3**. From this list of candidates, the top 12 compounds were selected for down-stream validation (**Figure 3**). Hits were classified into four distinct categories based upon their proposed mechanism of action, including (1) protein kinase and protease inhibitors, (2) anti-inflammatory compounds, (3) direct-acting antivirals, and (4) other host factor-targeting compounds. This set of hits contained compounds with previously reported SARS-CoV-2 antiviral activity (dinaciclib, GC376 sodium, cyclosporin A, and camostat mesylate) and seven compounds that have not previously been reported to have anti-SARS-CoV-2 activity **(Table 2) (*21–24*)**. These unreported compounds include the CDK inhibitor AZD5438, the AKT inhibitor SC66, the VEGFR inhibitor BFH772, the NADPH oxidase inhibitor GKT137831, the RAD51 inhibitor B02, the steroid budesonide, and the antiinflammatory compound cantharidin.

**Figure 3.**
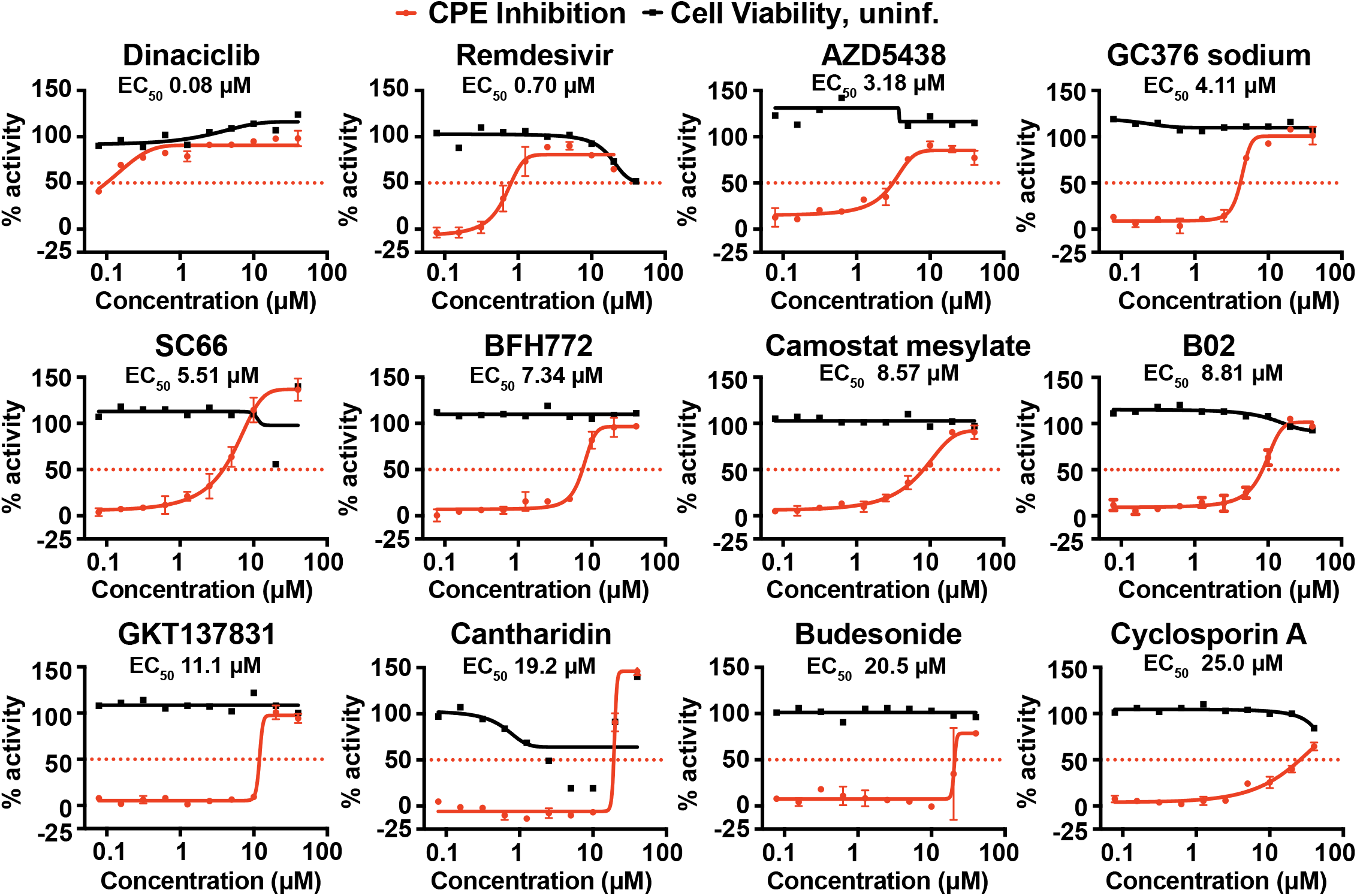
Dose response curves of compounds with SARS-CoV-2 antiviral activity. Calu-3 cells were infected with SARS-CoV-2 at MOI 0.05 and treated with compounds at indicated concentrations. Data show % CPE inhibition in SARS-CoV-2 infected cells (red) and % cell viability in uninfected cells (black). Data are normalized to the mean of DMSO-treated wells and represent mean ± SD for n = 2 technical replicates.

**Table 2.**

| Library | Compound | CellTiter Glo | TCID50 | qRT-PCR | IFA |  |  | Category | Previous reports | Clinical trial |
| --- | --- | --- | --- | --- | --- | --- | --- | --- | --- | --- |
|  |  | Calu-3 | Calu-3 | Calu-3 | Vero-E6 | HPMEC/hACE2 | Huh-7 |  |  |  |
| Bioactive | Dinaciclib | 0.08 | 0.13 | 0.08 | >10 | >10 | 0.04 | CDK inhibitor | Bouhaddou et al., Cell 2020, Ellinger et al., Research Square 2020, Wells et al., Nature |  |
| FDA | Remdesivir | 0.70 | 2.45 | 1.61 | 12.30 | 0.21 | > 40 | RDRP inhibitor | Jeon et al., BioRxiv 2020, Touret et al., BioRxiv 2020, Riva et al., Nature 2020 | yes |
| FDA | Camostat mesylate | 8.57 | 0.17 | 0.56 | >40 | >40 | >40 | TMPRSS2 inhibitor | Gordon et al., Nature 2020, Jeon et al., BioRxiv 2020, Vuong et al., Nature Communications | yes |
| Bioactive | AZD5438 | 3.18 | 4.62 | 2.41 | 5.25 | 10.28 | N.D. | CDK inhibitor | New |  |
| Bioactive | GC376 sodium | 4.11 | 10.16 | 5.22 | 1.61 | <1.25 | <1.25 | 3CLpro inhibitor | Ma et al., Cell Research 2020, Iketani et al., BioRxiv 2020 |  |
| Bioactive | Cantharidin | 19.20 | 4.84 | 1.05 | >40 | N.D. | N.D. | Anti-inflammatory | New |  |
| Bioactive | BFH772 | 7.34 | 5.78 | 24.92 | 19.45 | 5.80 | 4.80 | VEGFR kinase inhibitor | New |  |
| Bioactive | SC66 | 5.51 | 24.84 | 8.98 | N.D. | N.D. | N.D. | AKT Inhibitor | New |  |
| Bioactive | B02 | 8.81 | 20.35 | 18.24 | >40 | >40 | >40 | RAD51 inhibitor | New |  |
| Bioactive | GKT137831 | 11.10 | >40 | 10.13 | >40 | 17.02 | >40 | NADPH oxidase inhibitor | New |  |
| Bioactive | Apilimod mesylate | N.D. | 21.94 | 19.00 | 19.58 | <1.25 | <1.25 | Lipid kinase inhibitor | Bakowski et al., BioRxiv 2020, Dittmar et al., BioRxiv 2020, Riva et al., Nature 2020 | yes |
| FDA | Disulfiram | >40 | 15.17 | 13.16 | N.D. | 11.72 | >40 | 3CLpro inhibitor | Dittmar et al., BioRx 2020 | yes |
| FDA | Budesonide | 20.50 | 14.72 | >40 | >40 | 16.27 | >40 | Steroid | New | yes |
| FDA | Cyclosporin A | >40 | 7.54 | 36.72 | 14.41 | >40 | 2.78 | Calcineurin inhibitor | Dittmar et al., BioRxiv 2020, Jeon et al., BioRxiv 2020 | yes |

### Confirmation and characterization of SARS-CoV-2 antiviral candidate compounds

As our CPE-based screening assay measures SARS-CoV-2 induced cell death indirectly through quantification of ATP from viable cells, this assay does not directly test for antiviral activity of a given compound. To confirm the SARS-CoV-2 antiviral activity of our top 12 compound candidates, we tested their capacity to antagonize viral infection of Calu-3 cells through TCID50 and qRT-PCR assays. Also included was the PIKfyve kinase inhibitor apilimod mesylate, identified in Riva et al. 2020 (*22*). We found all tested candidates possessed significant antiviral activity with viral titer reductions of infectious virus and genome equivalents ranging from 1-4 logs compared to vehicle control treated cells **(Figure 4A-H)**. Intriguingly, we found that the CDK inhibitor dinaciclib (EC50 0.13 μM) was more potent in our assays than remdesivir (EC50 2.45 μM) in limiting viral titers in Calu-3 cells as measured by TCID50 and qRT-PCR **(Table 2)**.

**Figure 4.**
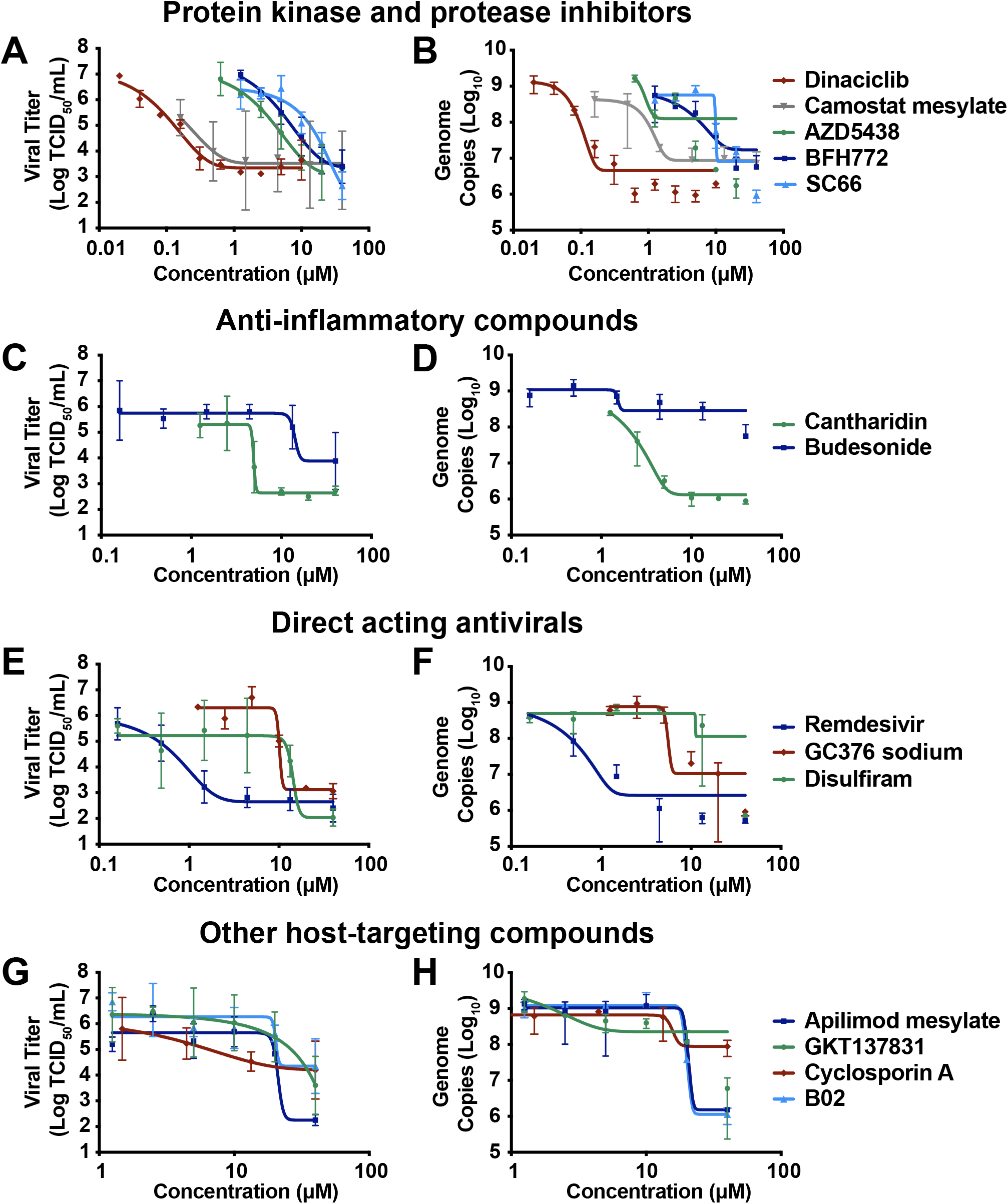
Confirmation and characterization of SARS-CoV-2 antiviral candidate compounds. Calu-3 cells were infected with SARS-CoV-2 at MOI 0.05, treated with the top 12 compounds (shown in Figure 3), disulfiram, or apilimod mesylate at indicated concentrations and supernatants were collected at 24 hpi. Viral titers and genome copies were calculated by TCID50 and qRT-PCR, respectively. **(A)** and **(B)** protein kinase and protease inhibitors, **(C)** and **(D)** anti-inflammatory compounds, **(E)** and **(F)** direct-acting antivirals and **(G)** and **(H)** other host-targeting compounds. TCID50 data represent mean ± SD for n = 2 independent experiments. Genome copy data represent mean ± SEM for n = 2 technical replicates and are representative of n = 2 independent experiments.

Since the antiviral activity of compounds *in vitro* may be cell-type specific, we tested the capacity of a smaller subset of our candidates (dinaciclib, camostat mesylate, BFH772, budesonide, GC376 sodium, apilimod mesylate, GKT137831, B02, and cyclosporin A) to antagonize virus infection across multiple cell types. Compounds were selected based on their lack of cytotoxicity across all cell lines (data not shown). We utilized an immunofluorescence confocal microscopy assay (IFA) to measure the capacity of these 8 compounds to inhibit SARS-CoV-2 replication in 3 diverse cell lines including Huh-7 (human hepatocytes), human pulmonary microvascular endothelial cells stably expressing the SARS-CoV-2 receptor human ACE2 (HPMEC/hACE2) (*25*), and Vero-E6 cells **(Table 1)**. Cells are treated with compound immediately prior to infection with SARS-CoV-2 and incubated for 24-48 hours. We then stained for the SARS-CoV-2 nucleoprotein (N) and calculated antiviral activity as % decrease in cell infection compared to vehicle treated infected cells. We found that the antiviral activity of GC376 sodium and apilimod mesylate were conserved across these cell lines **(Figure 5A-B)** while all other tested drugs exhibited cell type specific activity **(Figure 5C-I)**. GC376 sodium has been suggested to inhibit SARS-CoV-2 M protease (Mpro) (*26*), which may explain its efficacy across cell lines.

**Figure 5.**
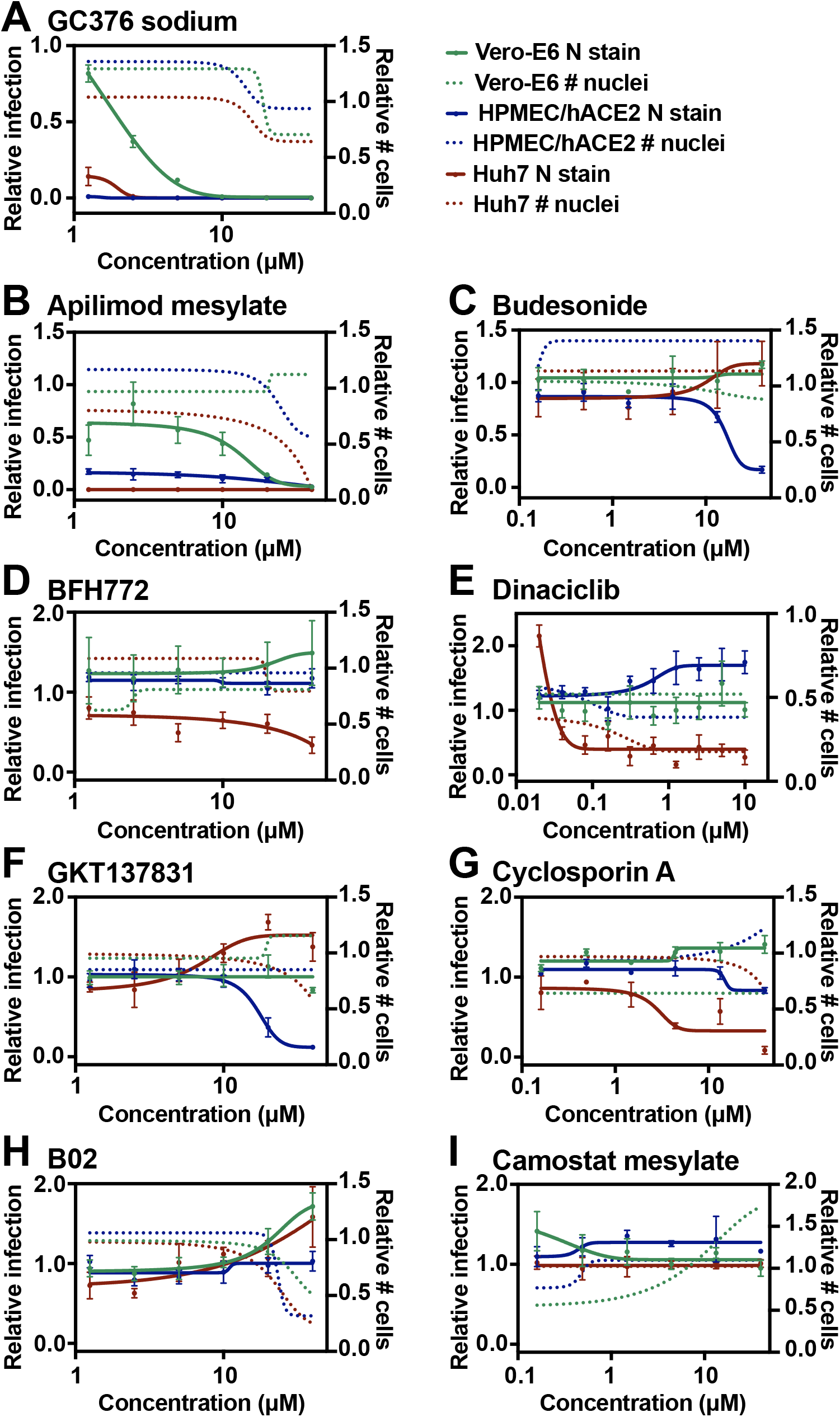
Cell-type specificity of compounds antiviral activity. Huh7, HPMEC/hACE2 and Vero-E6 cells were infected with SARS-CoV-2 at MOI 0.05 and treated with **(A)** Dinaciclib, **(B)** BFH772, **(C)** Budesonide, **(D)** GC376 sodium, **(E)** Apilimod mesylate, **(F)** GKT137831, **(G)** Cyclosporin A, **(H)** B02, and **(I)** Camostat mesylate at indicated concentrations. At 48hpi cells were washed, fixed, and stained with DAPI and for SARS-CoV-2 nucleocapsid protein. Plates were fluorescently imaged and analyzed for nucleocapsid stain per nuclei. Relative infection (full lines) and relative number of cells (dashed lines) are normalized to DMSO-treated wells. Data represent mean ± SEM for n = 4 technical replicates and are representative of n = 3 independent experiments.

To shed light on mechanisms of action, we tested whether our top compounds inhibit SARS-CoV-2 Mpro or papain-like protease (PLpro). Conducting in-house developed SARS-CoV-2 protease cleavage assays, we identified disulfiram as a potent PLpro inhibitor and both disulfiram and GC376 sodium as Mpro inhibitors, confirming previous observations (*26–28*) **(Figure S4)**. Notably, high concentrations of disulfiram exhibited antiviral activity in Calu-3 cells with 3 log reductions of viral titer and genome equivalents in TCID50 and qRT-PCR assays compared to vehicle control treated cells, potentially explaining its antiviral mechanism of action **(Table 2)**. In summary, these data highlight the benefit of a target-agnostic approach to expose previously unknown mechanisms of identified antivirals, as well as the importance of testing and validation across multiple cell lines for anti-viral screening.

### Candidate compound B02 exhibits antiviral synergy with the nucleoside analog remdesivir

Combination therapy is a highly desirable approach for treating SARS-CoV-2 infections (*29*). Because our screen revealed only a single compound (dinaciclib) possessing greater antiviral activity than remdesivir when used as a monotherapy, we asked if our compound candidates possess synergistic antiviral activity when used in combination with remdesivir. To assess this, we conducted a CPE inhibition assay using 10 μM of B02 in the presence or absence of an EC15 of remdesivir (2 μM, Figure S2B-C) in Vero-E6 cells. Interestingly, the RAD51 inhibitor B02 exhibited potent synergistic activity with remdesivir **(Figure 6A)**. We confirmed these results in Calu-3 cells by conducting a dose response of remdesivir in the presence or absence of 10 μM B02 and observed a >10-fold shift in the EC50 of remdesivir. from 1 μM to <0.08 μM (**Figure 6B**). Taken together, these observations suggest that in addition to providing new starting points for therapeutic development, B02 may be pursued as a combination therapy with remdesivir to develop treatments for COVID-19 patients.

**Figure 6.**
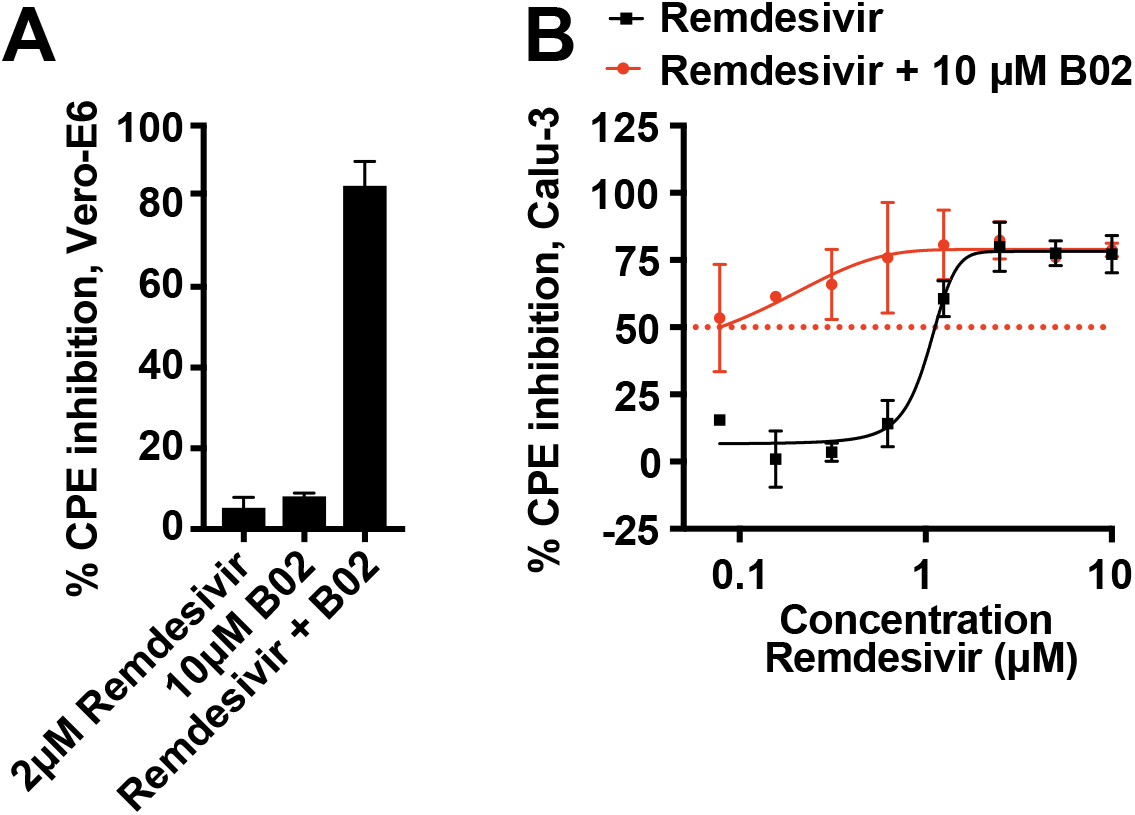
B02 synergy with remdesivir. **(A)** Vero-E6 cells were infected with SARS-CoV-2 at MOI 0.05 and treated with 2 μM remdesivir, 10 μM B02, or a combination of 2 μM remdesivir and 10 μM B02 for 72h. CPE inhibition was measured by CTG assay. **(B)** Calu-3 cells were infected with SARS-CoV-2 at MOI 0.05 and treated with remdesivir at indicated concentrations in the presence or absence of 10 μM B02 for 96h. CPE inhibition was measured by CTG assay and was normalized to DMSO-treated wells. Data represent mean ± SD for n = 2 technical replicates.

## Discussion

In this study, we screened a library of FDA-approved, clinical, and pre-clinical compounds for antiviral activity against SARS-CoV-2, with the goal of rapidly repurposing drug candidates for clinical use during the COVID-19 pandemic. We identified multiple cell lines that were permissive to SARS-CoV-2 infection and conducted primary screens using virus-induced CPE as a functional readout in two highly permissive but divergent cell types (Vero-E6 and Calu-3). We identified many compounds displaying varying degrees of antiviral activity across distinct cell lines including Vero-E6, Calu-3, HPMEC/hACE2, and Huh-7. These cell-type dependent phenotypes suggest significant cell type-dependency of compound efficacy *in vitro*. We further validated a subset of candidates to define EC50 values, confirmed antiviral activity in orthogonal assays, probed mechanism of action, and demonstrated antiviral synergy of one compound with remdesivir. While our investigation successfully identified compounds published previously, we also identified 7 potent compounds that to our knowledge have not been previously reported. They include the RAD51 inhibitor B02, the CDK inhibitor AZD5438, the AKT inhibitor SC66, the VEGFR inhibitor BFH772, the NADPH oxidase inhibitor GKT137831, the steroid budesonide, and the anti-inflammatory compound cantharidin. Further, B02 was found to synergize with remdesivir, shifting the apparent EC50 of remdesivir more than 10-fold. Together, our results present new possibilities for FDA-approved and other well-validated candidate drugs to be repurposed for remdesivir combination therapy.

The cell-type dependent antiviral efficacy of many of our drug candidates within this screen indicates a strong selection bias is introduced by the selection of cell type used in the screen. The biological reason for this is likely due to distinct cell-type expression patterns of host-factors essential for SARS-CoV-2 replication or differential drug metabolism. For example, SARS-CoV-2 requires proteolytic cleavage of the Spike (S) glycoprotein by a host protease once it binds to the ACE2 receptor (*30–32*). This cleavage can be performed by multiple host proteases including TMPRSS2 and cathepsin L, whose expression levels vary significantly between different cell lines (*30, 33, 34*). Vero-E6 cells do not express TMPRSS2, meaning any drug targeting TMPRSS2, or the interaction between S and TMPRSS2, would not emerge as a hit in such a cell line (*32*). Indeed the drug candidate camostat mesylate, a TMRPSS2 inhibitor currently in clinical trials (*35*) inhibited SARS-CoV-2 in Calu-3 but not Vero-E6 cells **(Figure 5)**. Conversely, GC376 sodium, a previously characterized SARS-CoV-2 protease inhibitor, maintains its antiviral activity across multiple cell lines in our study, suggesting that compounds targeting viral factors may be expected to show activity across different tissues, although the extent of activity can differ based on prodrug metabolism, as previously observed with remdesivir (*26, 27, 36*). Pharmacokinetics and dynamics must be taken into account for each compound and may be influenced by the metabolic state of distinct cell lines or tissue within infected humans. Our comparative cell line investigation of antiviral compounds highlights cell line selection as a critical step when conducting SARS-CoV-2 drug screens and may explain disparate data obtained in different studies (*21–23, 37–42*).

Defining the mechanisms of action of antiviral compounds is critical to determine how a compound may be modified to improve antiviral efficacy and to suggest the compounds’ pharmacokinetic and pharmacodynamic limitations. An advantage of screening libraries of well characterized compounds is that the molecular targets of many compounds are already determined. GSEA analysis of our top antiviral candidates revealed a set of enriched host and viral targets. Our candidate compounds possess distinct mechanisms of action including host kinase and protease inhibition, anti-inflammatory activity, and direct antiviral efficacy targeting virus factors. Our results also implicate host-pathways that are critical for SARS-CoV-2 viral replication including regulation of the cell cycle through modulating CDKs, regulating MAPK signaling through modulation of c-JUN N-terminal kinases (JNK), and modulation of glycogen synthase kinase 3 (GSK-3). Our screening results are consistent with other reports that these diverse cellular pathways are likely to be critical for the SARS-CoV-2 lifecycle (*21, 43–46*). These observations call for further mechanistic investigation of the dependency of SARS-CoV-2 on these various host pathways and highlight the potential of repurposing other drugs not studied here that target these pathways.

Though our study identified many distinct antiviral candidates, only dinaciclib exceeded the *in vitro* potency of remdesivir, suggesting that few, if any, “magic bullet” compounds exist in the library of FDA-approved and clinical/preclinical compounds. This is not unexpected for a repurposing approach, and therefore, combination therapy may be the best strategy to achieve high efficacy rapidly (*29*). Here, we identify the RAD51 inhibitor B02 as highly synergistic with remdesivir in both Vero-E6 and Calu-3 cells. The mechanism of action of this antiviral synergy is currently unclear and an active area of investigation. As RAD51 is known to play a role in DNA repair and strand exchange via interactions with CDK1 during the cell cycle, its mechanism of action may be connected to the dependency of SARS-CoV-2 on cell-cycle arrest during infection and repurposing of host machinery for viral replication (*21, 47–50*). Because remdesivir functions as a nucleoside analog that inhibits the function of the SARS-CoV-2 RNA-dependent RNA Polymerase (RdRP) during replication (*51, 52*), the antiviral activity of remdesivir may be enhanced when the virus is unable to modulate the cell cycle in such a way that promotes optimal viral RNA replication (*50*). Alternatively, RAD51 may directly promote SARS-CoV-2 replication as a component of the viral replication complex, as it is also reported to localize to the membranous replication complex of hepatitis c virus (HCV) where it interacts with the HCV nonstructural protein 3 (*53*). RAD51 may function comparably for SARS-CoV-2 and thus B02 may interfere with SARS-CoV-2 replication machinery in a manner that enhances the activity of remdesivir. This is further supported by the observation that B02 possesses antiviral activity in the absence of remdesivir suggesting either that RAD51 plays a direct role in promoting viral replication or that B02 has activity against other targets. We have previously observed antiviral synergy against SARS-CoV-2 for compounds that possess no antiviral activity on their own against SARS-CoV-2, such as the HCV NS5A inhibitor velpatasvir (*29*). This suggests antiviral synergy may arise through direct inhibition of one or more viral factors as well as through modulation of host pathways. Further, as remdesivir is a direct inhibitor of the RdRP, the virus can be thought of as in a “weakened” or “sensitized” state in the presence of remdesivir, potentially making it more vulnerable to additional chemical compounds with no appreciable activity on their own. Taken together, the potential of combinatorial approaches, ideally against distinct molecular targets, holds promise and should be prioritized for *in vivo* studies to determine efficacy.

At the beginning of the COVID-19 pandemic, many groups utilized a variety of genetic, proteomic, and chemical screening strategies to identify potential drugs for repurposing against SARS-CoV-2. These studies differ in the cell type selected for screening and in the assays used to determine viral replication. Most SARS-CoV-2 drug screens to date have utilized either an IFA-based approach or a CPE-based approach. In general, the IFA-based screens investigate antiviral activity earlier than CPE-based studies (typically 1-2 dpi vs. 3-4 dpi respectively) adding a selection bias for compounds that are effective at early time-points, as many compounds with mild antiviral activity (or shorter half-lives) may be overcome by viral replication at later time points. In addition to the need to interpret *in vitro* assay data carefully, our study highlights the importance of cell type selection when screening for antiviral compounds *in vitro*. Use of multiple cell lines is critical. In particular, many early studies used Vero cells due to their widespread availability, permissiveness to SARS-CoV-2 infection, and ease of use and manipulation. However, we found that activity of compounds against SARS-CoV-2 in Vero cells does not correlate well with activity in Calu-3 cells, an arguably more relevant human epithelial cell line. Thus, when interpreting *in vitro* antiviral candidates, both cell type, readout, and relative timing of drug and virus addition need to be carefully considered.

Although several COVID-19 vaccines have received Emergency Use Authorization recently by the FDA, pharmaceutical therapies for COVID-19 patients will continue to be urgently needed. Vaccines will not be widely available for several months, however more importantly, even an optimal vaccine will not eliminate severe COVID-19 due to limitations of vaccine use and efficacy in specific populations such as immunocompromised and pediatric patients. Our study identified 7 compounds not previously demonstrated to have antiviral activity with potential for COVID-19 therapy, one of them strongly synergistic with the approved COVID-19 therapeutic remdesivir. Intriguingly, budesonide is already studied in clinical trials for COVID-19 treatment because of its anti-inflammatory properties, and if positive, antiviral efficacy should be considered as an additional pharmacodynamic effect. Investigation into the mechanism of action and *in vivo* efficacy of these compounds is of paramount importance, as is defining the safety profiles of these compounds alone and in combination with remdesivir in humans. Such studies are currently underway in our lab as well as in other labs, but further collaboration is needed to expedite this process and provide access to pre-clinical and clinical testing to alleviate the global COVID-19 pandemic.

## Methods

### Cells lines

Multiple cell lines were acquired in this study to determine permissiveness to SARS-CoV-2 for use in drug screens. Vero-E6 cells were acquired from the American Type Culture Collection (ATCC) and maintained in DMEM media supplemented high glucose DMEM (Gibco, Waltham, MA) supplemented with 10% FBS (R&D Systems, Minneapolis, MN), 1X GlutaMAX (Gibco), and 1X PenStrep (Gibco) [D10 media]. Huh-7 and HPMEC cells were obtained from Dr. Eva Harris (UC Berkeley) and maintained in D10 media or endothelial growth medium 2 (EGM-2) using the EGM-2 bullet kit from Lonza, respectively. LNCaP, HaCaT, Caco-2, Calu-3, HCT-116, and A549 cells were obtained from the UC Berkeley Cell Culture Facility and maintained in D10 media. NCI-H1437 and RD cells were obtained from the Cell and Genome Engineering Core at UCSF via Dr. Olga Gulyaeva (UCSF) and Dr. Michael T. McManus (UCSF) and maintained in D10 media. HBEC-30KT and BEAS-2B cells were obtained from Dr. Neil Tay (UCSF) and Dr. Michael T. McManus (UCSF) via Dr. Patrick Mitchell (UC Berkeley) and Dr. Russell Vance (UC Berkeley) and maintained in defined keratinocyte serum free medium (catalog #10744019, ThermoFisher Scientific) and Advanced RPMI containing 5% FBS, 1% L-glutamine, 1X PenStrep, respectively. Huh-7.5.1 cells were obtained from Dr. Andreas Puschnik (Chan Zuckerberg Biohub) and maintained in D10 media. All cells were maintained in a C0_2_ incubator at 37° C with 5% CO_2_. HPMEC/hACE2 and A549/hACE2 cell lines were produced by transducing parental cells with lentivirus encoding the human ACE2 gene followed by puromycin selection (2 μg/ml) for three passages. The hACE2 encoding plasmid was a gift from Hyeryun Choe (Addgene plasmid # 1786; htt://n2t.net/addgene:1786; RRID:Addgene_1786) (*54*).

### SARS-CoV-2 stock preparation and infections

The USA-WA1/2020 strain of SARS-CoV-2 was obtained from BEI Resources. The initial stock from BEI was passed through a 0.45 μM syringe filter and 5 μl of this filtered stock was inoculated onto ~80% confluent T175 flasks (Nunc, Roskilde, Denmark) of Vero-E6 cells to produce passage 1 of the virus. CPE was monitored daily and flasks were frozen down when cells exhibited 50-70% cytopathic effect (CPE) (48-72 hpi). Thawed lysates were then collected and cell debris was pelleted at 3000 x rpm for 20 minutes. Clarified viral supernatant was then aliquoted and infectious virus was quantified via a TCID50 assay. To produce passage 2 SARS-CoV-2 working stocks, 5 μl of the passage 1 stock was inoculated onto ~80% confluent T175 flasks of Vero-E6 cells as described above. Viral titers produced in Vero-E6 cells ranged from 1×10^6^-5×10^6^ TCID50 units / mL.

### Compound preparation, drug screening, and synergy experiments

The compound screening and 384-well infection experiments were conducted as described previously (*29*) and are described below. The FDA-approved drug library (Targetmol) and the Bioactives Library (Targetmol, Wellesley Hills, MA) containing 1,200 compounds and 4,170 compounds, respectively, were stored in dimethyl sulfoxide (DMSO) at 10 mM in 384-well master plates. Remdesivir (T7766, Targetmol) was also stored at 10 mM in DMSO. For drug screening, 2.5×10^3^ Vero-E6 cells (12 μl/well) or 1×10^4^ Calu-3 (12 μl/well) were seeded in 384-well white optical-bottom tissue culture-treated plates (Nunc) with a Multidrop Combi liquid handling instrument (Thermo Fisher Scientific, Waltham, MA). Cells were incubated for 24 (Vero-E6) or 48 hours (Calu-3) at 37°C and 5% CO_2_ before experiments were conducted. For the primary screen, dose response experiments, and synergy experiments, compounds were prediluted to 4x final concentration (8x for synergy experiments) in high glucose DMEM. 6 μl and 3 μl of media (for primary and synergy experiments, respectively) was transferred from compound dilution plates to cells in 384-well plates using a Cybio Well vario liquid handler (Analytik Jena, Jena, Germany), leading to a final concentration of DMSO at 0.4% (v/v) in the assay plate. Primary screens were conducted at 40 μM compound. For the dose response experiments, 10-point dose responses were generated by conducting 2-fold dilutions starting at 40 μM for compound confirmation and 10 uM for remdesivir in synergy plates. For all experiments conducted above, cells were incubated with compounds at 37°C and 5% CO_2_ for 1 hour before infection.

For all experiments above, cells were infected in 384-well plates at a multiplicity of infection (MOI) of 0.05 in a total volume of 6 μl/well. Cells were harvested for CTG-analysis once complete CPE was observed in DMSO-treated infected wells (72 hpi for Vero-E6 and 96 hpi for Calu-3). For harvest, opaque stickers (Nunc) were applied to the bottoms of plates to minimize signal spillover between wells, and plates were developed with the CellTiter-Glo 2.0 reagent (Promega, Madison, WI) according to the manufacturer’s instructions, with the exception of Vero-E6 cells, for which CTG reagent was diluted 1:1 (v/v) in PBS (Gibco, Waltham, MA, USA). Luminescence was read on a Spectramax L (Molecular Devices, San Jose, CA). Each plate contained 24 wells of uninfected/DMSO treated cells (100% CPE inhibition), and 24 wells infected/DMSO treated cells (0% CPE inhibition). Average values from those wells were used for data normalization and to determine % CPE inhibition for test compound wells. Duplicate plates were used to calculate average values and standard deviations. Z’ was determined as described previously (*55*). Statistical significance between experimental conditions were assessed using a two-tailed, heteroscedastic student’s t-test. Measurements were taken from distinct samples unless indicated otherwise. The data was plotted and analyzed with Spotfire (Tibco) and GraphPad Prism (San Diego, CA).

### GSEA Analysis

The methods for GSEA analysis conducted for this study were previously reported and described below (*29*). In brief, candidate compounds identified in our screens were assigned distinct properties based on known host targets and pathways using the Center for Emerging and Neglected Diseases’ database and for pharmacokinetic data and transporter inhibition data, the DrugBank database. Each assigned property was tested for enrichment among the screening hits using the gene set enrichment analysis (GSEA) software as described (*22, 56, 57*). Compounds annotated for each property were considered as part of the “gene set”. For each set of annotations, the background compound set was defined as the set of compounds annotated for any property. GSEA preranked analysis was performed using the compounds’ % CPE inhibition from each screen. Compound sets included in the analysis were between 5 and 500 compounds. The enrichment score (ES) reflects the degree to which a gene set is overrepresented at the top of a ranked list of compounds interacting with the given target. GSEA calculates the ES by walking down the ranked list of compounds interacting with the given target, increasing a running-sum statistic when a gene is in the gene set and decreasing it when it is not.

### Immunofluorescence microscopy analysis (IFA)

1×10^4^ Vero-E6, HPMEC/hACE2, or Huh-7 cells were seeded in black 96-well plates with clear bottoms 24 hours before adding drug combinations and infecting with SARS-CoV-2 at MOI 0.05 (viral inoculums were not washed away). Plates were fixed in 4% paraformaldehyde (PFA) 24 hpi (Vero-E6 and HPMEC/hACE2) and 48 hpi (Huh-7), permeabilized using 0.2% saponin in blocking buffer (2% BSA, 1% FBS in PBS) at room temperature for 30 minutes, incubated with mouse anti-SARS-CoV-2 nucleocapsid protein (1:1000, Sino Biological, Beijing, China; 40145-MM05) overnight in blocking buffer, incubated with goat anti-mouse AlexaFluor647 (1:1000, Abcam, Cambridge, United Kingdom) and DAPI/Hoechst (1:1000, Invitrogen) in blocking buffer, fixed in 4% PFA, and kept in 1x PBS until imaging on an Image Xpress Micro 4 (Molecular Devices). An average of 1×10^3^ cells were imaged across four sites per well and were analyzed for nucleocapsid (N) stain per nuclei (DAPI) using CellProfiler 3.1.9 (Broad Institute, Cambridge, MA).

### TCID50 assay

5×10^4^ Calu-3 cells were seeded into 96-well plates 48 hours before adding drug combinations and viral inoculum (MOI 0.05) (viral inoculum was not washed away). At 24 hpi supernatant was collected from each well and serially diluted, and each dilution was applied to eight wells in 96-well plates containing Vero-E6 cells. Three days later, CPE was counted visually and TCID50/mL was calculated using the dilution factor required to produce CPE in half, or 4/8, of the wells for a given dilution.

### RT-qPCR

RT-qPCR was conducted as previously reported (*29*), and further described below. For RT-qPCR, supernatants were collected at 48hpi and inactivated 1:1 in 1X DNA/RNA Shield for RNA extraction and RT-qPCR analysis (Zymo Research, Irvine, CA). RNA was extracted using the QIAamp Viral RNA Mini Kit (Qiagen, Hilden, Germany) according to the manufacturer’s instructions. In brief, 140 μL of each sample was mixed with 560 μL of Carrier-RNA-containing AVL and incubated for 10 min at RT. After addition of 560 μL of 100% ethanol, the samples were spun through columns. The columns were washed sequentially with 500 μL of AW1 and 500 μL AW2 and RNA was eluted using 50 μL of RNAse free water. RT-qPCR reactions with TaqPath master mix (Thermo Fisher) were assembled following the manufacturer’s instructions. For a 20 μL reaction, 5 μL of 4x TaqPath master mix was combined with 1.5 μL of SARS-CoV-2 (2019-nCoV) CDC N1, N2, or RNase P qPCR Probe mixture (Integrated DNA Technologies, Cat. #10006606, Primer sequences: 2019-nCoV_N1-F 2019-nCoV_N1: GAC CCC AAA ATC AGC GAA AT; 2019-nCoV_N1-R 2019-nCoV_N1: TCT GGT TAC TGC CAG TTG AAT CTG; 2019-nCoV_N1-P 2019-nCoV_N1 FAM: ACC CCG CAT TAC GTT TGG TGG ACC-BHQ1), RNA sample, and water to a final volume of 20 μL. Volumes were divided by 2 for 10 μL reactions. RT-qPCR was performed on a BioRad CFX96 or CFX384 instrument with the following cycle: 1) 25°C for 1min, 2) 50°C for 15min, 3) 95°C for 2min, 4) 95°C for 3s, 5) 55°C for 30s (read fluorescence), 6) go to step 4 for 44 repetitions. Quantification cycle (Cq) values were determined using the second derivative peak method (*58*). Custom code written in MATLAB (available at https://gitlab.com/tjian-darzacq-lab/second-derivative-cq-analysis) was used to take the numerical second derivative of fluorescence intensity with respect to cycle number, using a sliding window of ± 3 cycles. The peak of the second derivative was fit to a parabola, whose center was taken to be the Cq value (*58*).

### SARS-CoV-2 MPro Activity Assay

Compounds were dissolved in DMSO at 50X the desired screening concentration. DMSO was used as a solvent control. MPro protein (purification described below) was diluted in assay buffer (50 mM HEPES, pH 7.5, 150 mM NaCl, 1 mM EDTA, 0,01% pluronic acid F127) to a concentration of 30 nM and 24.5 μL of diluted protein was aliquoted to each well of a black 384 well plate (Corning 384-Well, Flat-Bottom Microplate). Each well was treated with 0.5 μL of compound or vehicle and the plate was incubated for 30 min at room temperature. During the compound incubation the peptide probe KTSAVLQ-Rh110-gammaGlu (Biosyntan) was diluted from 5 mM DMSO stock into assay buffer. After pre-incubation, 5 μL of 75 μM Rh-110 probe was added to each well. RFU value was immediately measured on a Tecan Spark plate reader with an excitation wavelength of 488 nm and an emission wavelength of 535 nm at 30 °C for 30 min.

### Purification of SARS-CoV-2 Main Protease

The coding sequence for SARS-CoV-2 main protease was codon-optimized for E. coli and synthesized by Integrated DNA Technologies. The sequence was amplified by PCR and cloned into the pGEX6P-1 vector, downstream of GST and an HRV 3C protease cleavage site, using the Gibson Assembly Master Mix kit (New England BioLabs, Inc). To ensure authentic termini, the amino acids AVLQ were added to the N-terminus of the main protease by addition of their coding sequence to the 5’ end of the gene product. This sequence reconstitutes the SARS-CoV-2 NSP4/5 cleavage site, resulting in auto-cleavage by the main protease protein product (*59*). Similarly, we added a GP-6xHis tag for IMAC purification to the C-terminus (the GP completes a non-consensus 3C cleavage site along with the C-terminus of the main protease which allows for cleavage of the his tag after purification, resulting in an authentic C-terminus). Hi Control BL21(DE3) cells were transformed with the expression plasmid using standard techniques. We used Hi Control cells as we observed expression of the main protease was toxic in other standard E. coli cell lines. A single colony was used to start an overnight culture in LB + carbenicillin media. This culture was used to inoculate 2 x 1 L cultures in Terrific Broth, supplemented with 50 mM sodium phosphate pH 7.0 and 100 μg/mL carbenicillin. These cultures grew in Fernbach flasks at 37 °C while shaking at 225 rpm, until the OD600 reached approximately 2.0, at which point the temperature was reduced to 20 °C and 0.5 mM IPTG (final) was added to each culture. The cells were allowed to grow overnight.

The next day, the cultures were centrifuged at 6,000 x g for 20 minutes at 4 °C, and the resulting cell pellets were resuspended in IMAC_A buffer (50 mM Tris pH 8.0, 400 mM NaCl, 1 mM TCEP). Cells were lysed with two passes through a cell homogenizer (Microfluidics model M-110P) at 18,000 psi. The lysate was clarified with centrifugation at 42,000 x g for 30 minutes and the cleared lysate was loaded onto 3 x 5 mL HiTrap Ni-NTA columns (GE) pre-equilibrated with IMAC_A buffer, using an AKTA Pure FPLC. After loading, the columns were washed with IMAC_A buffer until the A280 levels reached a sustained baseline. The protein was then eluted with a linear gradient with IMAC_B buffer (50 mM Tris pH 8.0, 400 mM NaCl, 500 mM imidazole, 1 mM TCEP) across 25 column volumes, while 2 mL fractions were collected automatically. Peak fractions were analyzed by SDS-PAGE and those containing SARS-CoV-2 main protease were pooled. Importantly, auto-cleavage of the N-terminal GST tag was observed and the eluted protein had a mass consistent with SARS-CoV-2 main protease along with the C-terminal GP-6xHis tag, as determined by ESI-LC/MS.

Pooled fractions were treated with HRV 3C protease (also known as “PreScission” protease) while dialyzing against IMAC_A buffer at room temperature (2 x 2 L dialyses). Room temperature dialysis was important as we observed a tendency for the main protease protein to precipitate with prolonged exposure to 4 °C. Cleavage of the C-terminal GP-6xHis tag was confirmed after 2 hours by ESI-LC/MS. The dialyzed and cleaved protein was then re-run through a 5 mL HiTrap Ni-NTA column pre-equilibrated with IMAC_A buffer. The main protease eluted in the flow-through as expected.

The protein was then concentrated to approximately 5 mL and loaded onto a Superdex 75 16/60 column pre-equilibrated with SEC Buffer (25 mM HEPES pH 7.5, 150 mM NaCl, 1 mM TCEP). The protein was run through the column at 1 mL/min and eluted as one large peak well in the included volume (at ~75 mL). Fractions from this peak were analyzed by SDS-PAGE and pure fractions were pooled and concentrated to 10 mg/mL, aliquoted, and stored at −80 °C. Final yield was typically in the realm of 60-70 mg/L of culture.

### SARS-CoV-2 PLpro Activity Assay

Compounds were dissolved in DMSO at 50X the desired screening concentration. DMSO was used as a solvent control. PLpro protein was purified as described below. The screening assay was performed in black 384 well plates (Corning 384-Well, Flat-Bottom Microplate), and was performed in a 25.5 μL volume which contained a final PLpro concentration of 50 nM, a 50 μM concentration of substrate (RLRGG-AMC), and 0.5μl of DMSO or compound (final concentration of 40μM), the final assay buffer contained 20mM Hepes pH 7.5, 100mM NaCl and 0.1% mg/ml BSA. Screens were performed with 1:5000 antifoam to reduce the surface tension and bubbles. After addition of the substrate, the RFU value was measured on a Tecan Spark plate reader with an excitation wavelength of 360 nm and an emission wavelength of 460 nm at 30 °C for 30 min.

### Expression and purification of PLpro

The papain-like protease (PLpro) expressing plasmid, 2BT-Nsp3-PLpro was transformed into E. coli BL21 (DE3) and plated on ampicillin resistant LB agar plate. The next day, a colony was picked up for overnight culture in the presence of ampicillin 100μg/ml. For large-scale protein purification, a 1L culture of 2XYT media was grown using overnight culture (1:100) at 37°C (210 rpm). The bacterial culture was grown to an OD600 ~0.8-1.0 and induced with 1mM IPTG. The protein was expressed at 20°C for overnight (18-20 hours). The bacterial culture was harvested at 4000 X g and cell pellets were resuspended in 30 mL lysis buffer (25mM Tris-HCl pH 8.0, 250mM NaCl, 10% glycerol, 5mM β-mercaptoethanol), supplemented with protease inhibitor tablets. The cell culture was sonicated at 20% amplitude for 7 minutes (0.5 sec ON, 1.5 sec OFF). Cellular debris was pelleted down by centrifuging at 15,000 X g for 20 minutes at 4°C. The supernatant was loaded on a Talon column (GE Healthcare Life Sciences) (pre-equilibrated with lysis buffer) at a speed of 1 mL/min. Non-specific proteins were washed with 20 column volumes of Buffer-A (lysis buffer supplemented with 25 mM imidazole). PLpro protein was eluted with 5 column volumes of Buffer-B (lysis buffer supplemented with 250 mM imidazole). The eluted protein was concentrated using a 10 kDa MWCO filter (Amicon-Millipore), and concentrated up to 2 mg/mL.

## Acknowledgements

We thank Dr. Ella Hartenian (UC Berkeley) and Dr. Britt Glaunsinger (UC Berkeley) for assistance with cloning the human ACE2 gene into a lentivirus construct. We thank Dr. Patrick S. Mitchell (UC Berkeley/University of Washington), Dr. Olga Gulyaeva (UCSF), Dr. Andreas Puschnik (Chan Zuckerberg Biohub), and Dr. Eva Harris for cell lines used in this study and helpful discussion. We further thank members of the Harris and Stanley labs for helpful discussion. We would like to acknowledge receipt of diverse cell lines used in this study from the UCB Cell Culture facility, which is supported by the University of California, Berkeley. SARS-CoV-2 clinical isolate USA-WA1/2020, NR-52281 was obtained through BEI resources, NIAID, NIH and deposited by the Centers of Disease Control and Prevention. This project was supported by generous funding from Eric and Wendy Schmidt by recommendation of the Schmidt Futures program, through Covid Catalyst funding administered by the Center of Emerging and Neglected Diseases at UC Berkeley, and through Fast Grants (part of Emergent Ventures at George Mason University). SBB is an Open Philanthropy Awardee of the Life Science Research Foundation. EVD is supported by NSF Graduate Research Fellowship DGE-1752814. CDD and TGWG are supported by the Bowes Research Fellows Program N7342, the Siebel Stem Cell Institute W6188, the Jane Coffin Childs Memorial Fund for Medical Research, and the HHMI.

## Author Contributions

Conceptualization: S.B.B., E.V.D., E.W., L.H.Y, X.N, Do.M.F, J.S, S.A.S.

Funding acquisition: M.O., N.M, D.K.N., J.S., S.A.S.

Investigation: S.B.B., E.V.D., E.W., L.H.Y., X.N., C.D., T.G.W.G., J.R.S., G.R.G., A.W.R., Da.M.F., J.N.S., C.C.W., T.B., D.D., U.S.G., R.B., Do.M.F.

Methodology: S.B.B., E.V.D., E.W., L.H.Y., X.N., N.M., D.K.N., J.S., S.A.S.

Project administration: J.S., S.A.S.

Supervision: M.O., N.M., D.K.M., J.S., S.A.S.

Visualization: S.B.B., E.V.D., E.W., L.H.Y, X.N, J.R.S., Do.M.F, J.S, S.A.S.

Writing-original draft: S.B.B.

Writing-review and editing: S.B.B., E.V.D., E.W., L.H.Y, X.N, Do.M.F, J.S., S.A.S.

---

**Figure S1.**
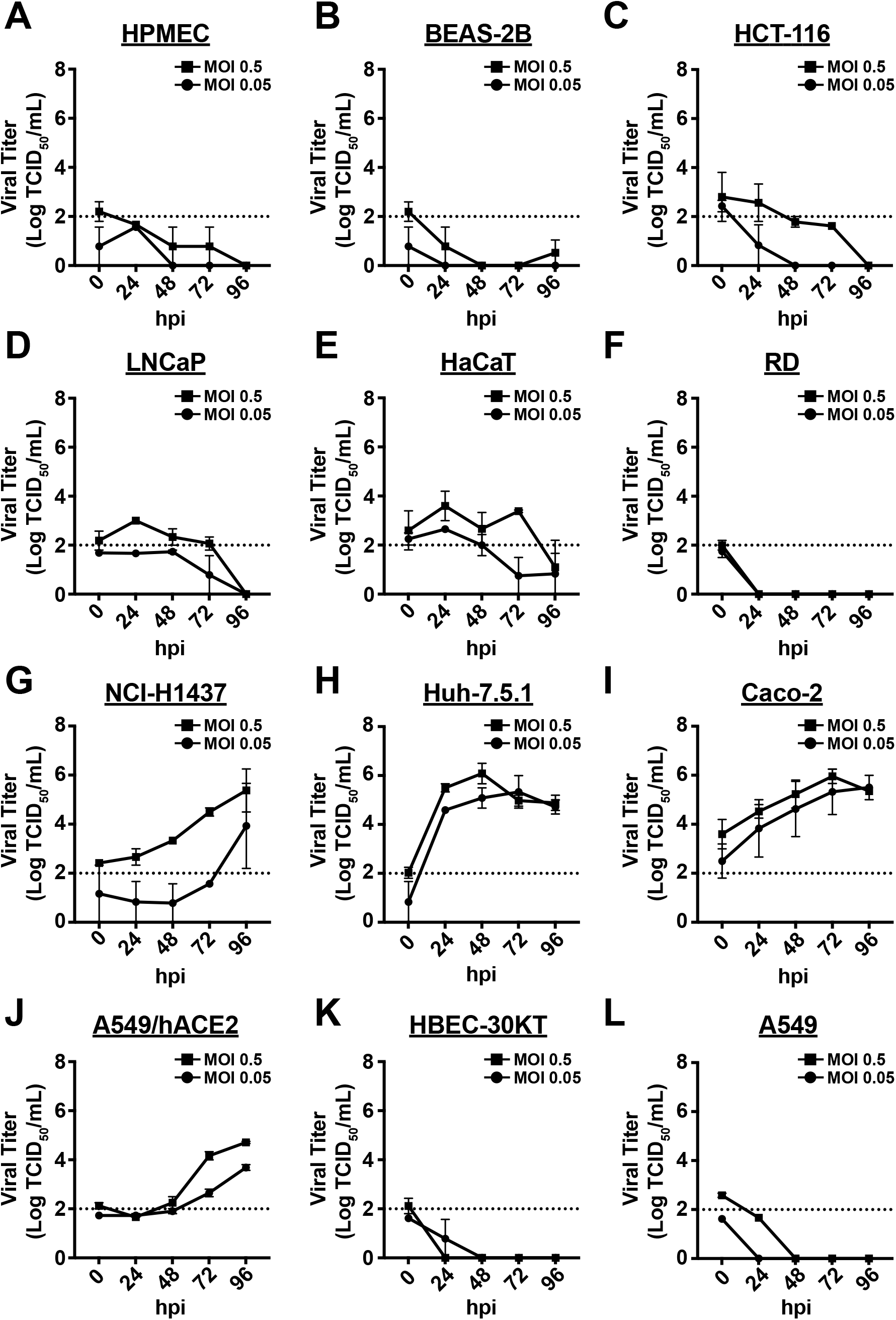
**(A)** HPMEC, **(B)** BEAS-2B, **(C)** HCT-116, **(D)** LNCaP, **(E)** HaCaT, **(F)** RD, **(G)** NCI-H1437, **(H)** Huh-7.5.1, **(I)** Caco-2, **(J)** A549/hACE2, **(K)** HBEC-30KT, or **(L)** A549 cells were infected with SARS-CoV-2 at MOI 0.5 or 0.05 as in figure 1. Viral titers were analyzed by TCID50 assay at the indicated time points (hours post-infection, hpi). Dashed lines represent limit of detection. Data represent mean ± SEM for n = 2 independent experiments.

**Figure S2.**
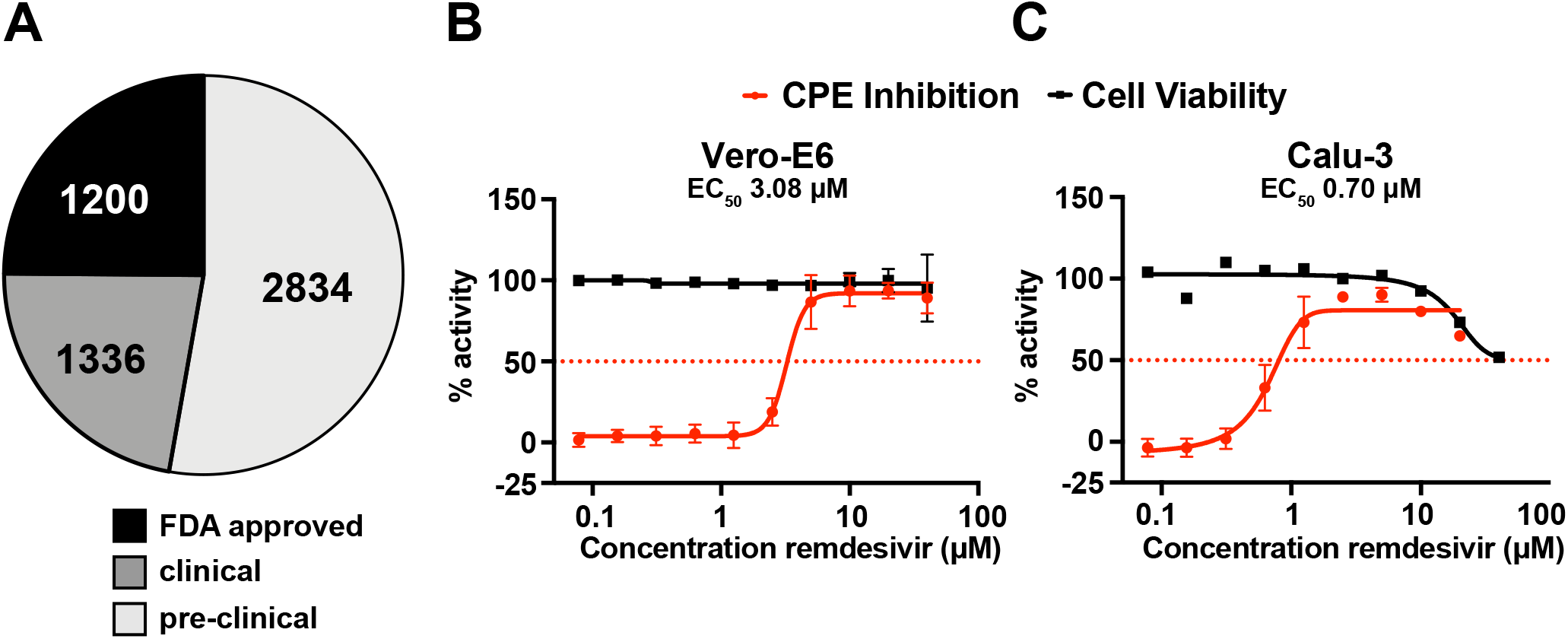
**(A)** Clinical status of compounds tested. **(B-C)** Remdesivir dose response curves in **(B)** Vero-E6 and **(C)** Calu-3 cells showing % CPE inhibition in SARS-CoV-2 infected cells (red) and % cell viability in uninfected cells (black). Data are normalized to the mean of DMSO-treated wells and represent mean ± SD for n = 2 technical replicates.

**Figure S3.**
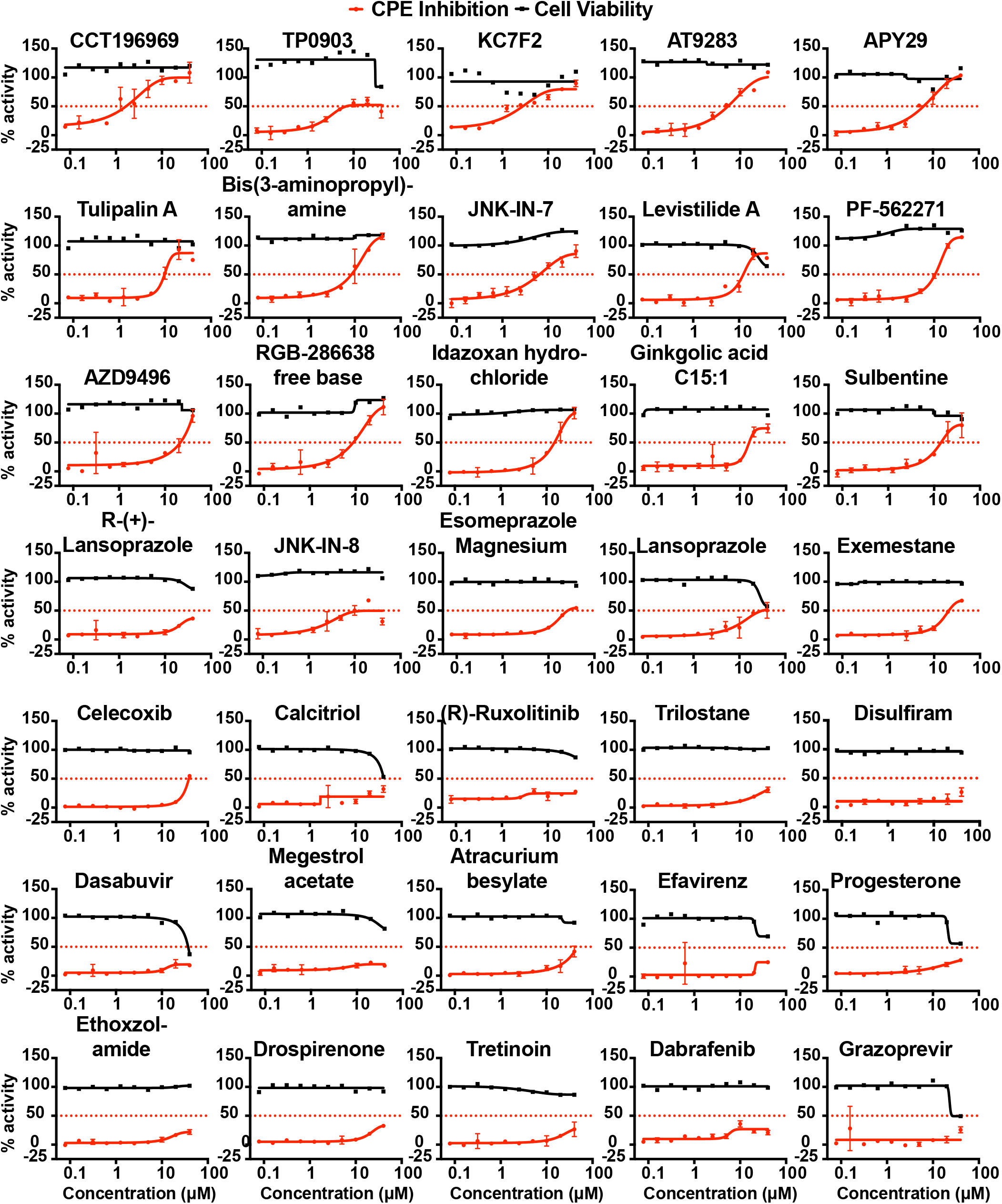
Calu-3 cells were infected with SARS-CoV-2 at MOI 0.05 and treated with compounds at indicated concentrations. Data show % CPE inhibition in SARS-CoV-2 infected cells (red) and % cell viability in uninfected cells (black). Data are normalized to the mean of DMSO-treated wells and represent mean ± SD for n = 2 technical replicates.

**Figure S4.**
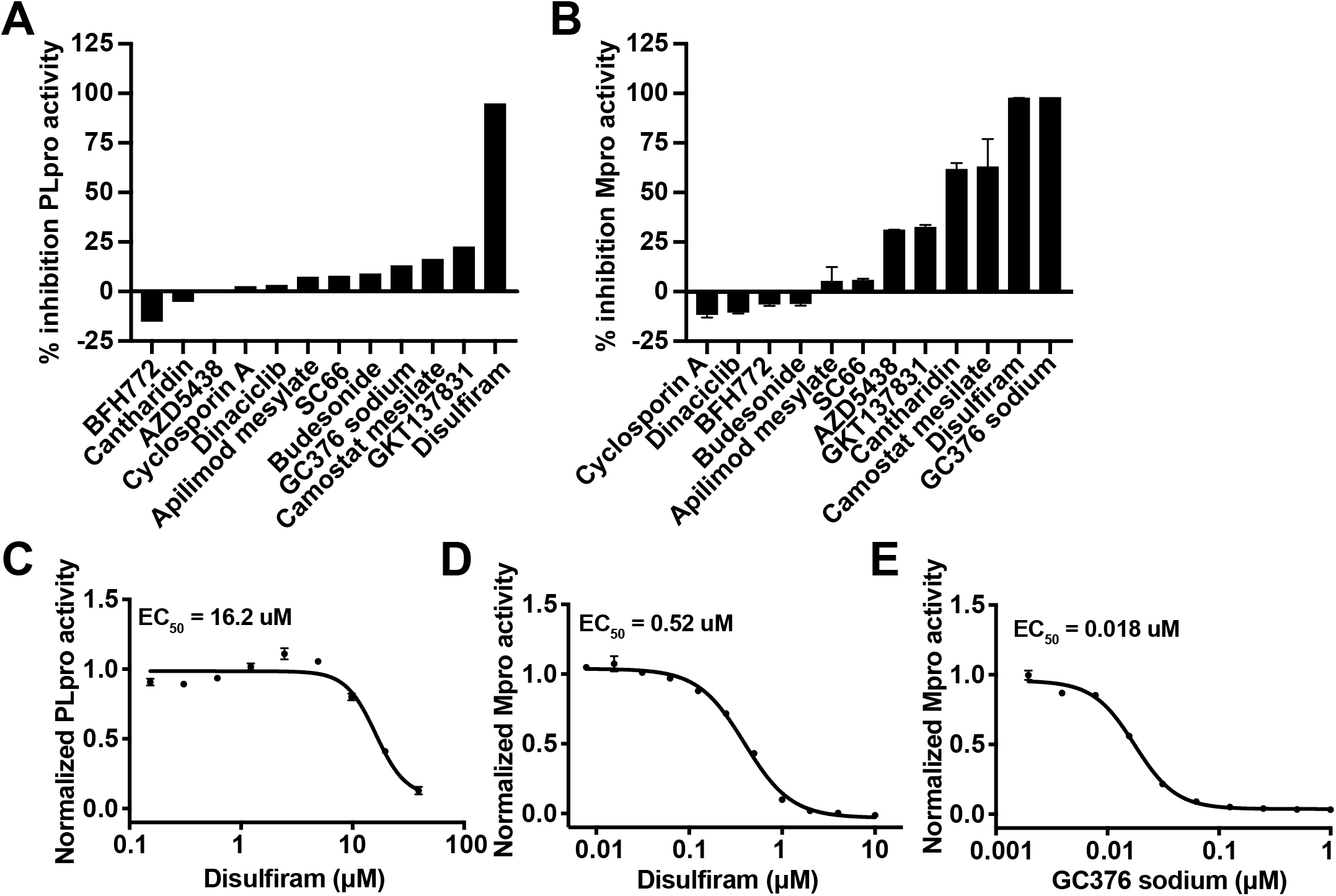
**(A-B)** Percent inhibition of of **(A)** PLpro or **(B)** Mpro activity was measured. **(C-D)** Disulfiram dose response curve of inhibition of **(C)** PLpro and **(D)** Mpro activity. **(E)** GC376 sodium dose response curve of inhibition of Mpro activity. Data represent n = 1 (A) and mean ± SEM for n = 3 technical replicates (B-E).

